# An alternative tetrahydrofolate pathway for formaldehyde oxidation in verrucomicrobial methanotrophs: Primer design for *folD* and *ftfL* and transformation of *E. coli*

**DOI:** 10.1101/2022.08.24.504922

**Authors:** Rob A. Schmitz, Koen A.J. Pelsma, Huub J.M. Op den Camp

## Abstract

Methylotrophs make a living by using one-carbon compounds as energy and carbon source. Methanol (CH_3_OH) is utilized by various methylotrophs and is oxidized by a methanol dehydrogenase. The calcium-dependent methanol dehydrogenase MxaFI converts methanol to formaldehyde (CH_2_O). In addition to MxaFI, the lanthanide-dependent methanol dehydrogenase XoxF is found in a wide range of bacteria. XoxF isolated from the verrucomicrobial methanotroph *Methylacidiphilum fumariolicum* SolV possesses an unusually high affinity for both methanol and formaldehyde and converts methanol to formate (HCOOH) *in vitro*. However, genomic analyses and biochemical studies on related XoxF methanol dehydrogenases have questioned whether these enzymes are dedicated to the conversion of formaldehyde to formate *in vivo*. Instead, the genes encoding the bifunctional enzyme FolD and the enzyme FtfL, which we detected in all verrucomicrobial methanotrophs, were proposed to form a formaldehyde oxidation pathway utilizing tetrahydrofolate as C1-carrier. *folD* and *ftfL* are expressed in *M. fumariolicum* SolV and most closely related to homologues found in the phyla Verrucomicrobia and Proteobacteria, respectively. Here, we designed primers targeting Mf-*folD* and Mf-*ftfL* and amplified these genes through PCR. The amplified genes were ligated into pET30a(+) vectors which were subsequently used for the successful transformation of *E. coli* XL1-Blue cells. The vectors can be used for heterologous expression and subsequent His-tag purification to biochemically investigate whether FolD and FtfL could form an alternative tetrahydrofolate pathway for formaldehyde oxidation in verrucomicrobial methanotrophs.

## INTRODUCTION

Methylotrophs utilize one-carbon molecules such methanol (CH_3_OH) as a source of energy and carbon (Chistoserdova and Kalyuzhnaya, 2018). Methanotrophs are a unique type of methylotrophs that make a living from the oxidation of methane (CH_4_). Aerobic methanotrophs are either members of the phylum Verrucomicrobia or the phylum Proteobacteria (Hanson and Hanson, 1996; Op den Camp et al., 2009). Methane oxidation is initiated by the conversion of methane to methanol via a soluble and/or particulate methane monooxygenase (Ross and Rosenzweig, 2017). Subsequently, methanol is oxidized by the calcium-dependent pyrroloquinoline quinone (PQQ) methanol dehydrogenase (MDH) MxaFI or the lanthanide-dependent PQQ-MDH XoxF (Keltjens et al., 2014; Picone and Op den Camp, 2019). Although XoxF was only recently discovered, it is becoming increasingly clear that XoxF-type MDHs are more diverse and widespread compared to MxaFI-type MDHs (Pol et al., 2014; Picone and Op den Camp, 2019). Additionally, MxaFI is hypothesized to have evolved from a XoxF prototype (Keltjens et al., 2014). Whereas MxaFI produces formaldehyde (CH_2_O) from the oxidation of methanol, XoxF isolated from the verrucomicrobial methanotroph *Methylacidiphilum fumariolicum* SolV was shown to produce formate (CHOOH) *in vitro* (Pol et al., 2014). The synthesis of formate as product could be attributed to the unusually high affinity of XoxF for both methanol and formaldehyde in comparison to MxaFI. However, a XoxF homologue of the proteobacterial methylotroph *Methylobacterium extorquens* AM1 was shown to produce formaldehyde *in vivo*, questioning the product released by XoxF-type MDHs (Good et al., 2019).

Formaldehyde is a central intermediate in many methylotrophs that assimilate this carbon source via the ribulose monophosphate (RuMP) cycle or the serine cycle (Chistoserdova, 2011). Verrucomicrobial methanotrophs lack a complete RuMP or serine cycle (Op den Camp et al., 2009). Instead, these bacteria were shown to use the Calvin-Benson-Bassham (CBB) cycle for carbon fixation, which incorporates carbon at the level of CO_2_ (Khadem et al., 2011). In verrucomicrobial methanotrophs assimilation is therefore unlikely to occur at the level of formaldehyde. Still, CO_2_ is produced from the oxidation of formaldehyde, but the route from formaldehyde to CO_2_ is not fully understood. Hou et al. (2008) hypothesized that the verrucomicrobial methanotroph *Methylacidiphilum infernorum* V4 could use a pathway in which the cofactor tetrahydrofolate (H_4_F) is involved as C1-carrier. Within this pathway, enzymes encoded by the genes *folD* and *ftfL* were proposed to convert formaldehyde to formate via the intermediates methylene-tetrahydrofolate (CH_2_-H_4_F), methenyl-tetrahydrofolate (CH-H_4_F) and formyl-tetrahydrofolate (CHO-H_4_F). The genes *folD* and *ftfL* are present in a large variety of organisms for the conversion of one-carbon compounds (Vorholt, 2002). Comparison of the genomes of eleven verrucomicrobial methanotrophs revealed that they all possess *folD* and *ftfL* (Schmitz et al., 2021; Picone et al., 2021). The oxidation of formaldehyde to formate via the enzymes FolD and FtfL could result in the direct production of NAD(P)H and ATP (Goenrich et al., 2002; Marx et al., 2003). In contrast, when instead XoxF would convert methanol to formate, two extra electrons would be released and presumably transferred to a terminal oxidase (Good et al., 2019). A separate formaldehyde oxidation pathway is therefore favourable in terms of energy conservation (Keltjens et al., 2014).

To investigate the involvement of FolD and FtfL in the conversion of formaldehyde to formate in verrucomicrobial methanotrophs, purification of these enzymes is necessary. Accordingly, primers were designed for the amplification of *folD* and *ftfL* of *M. fumariolicum*SolV (*Mf-folD* and *Mf-ftfL*, respectively). Here, we show the successful usage of these primers for the amplification of the genes of interest. The PCR products were ligated into pET30a(+) vectors and *E. coli* XL-1 Blue cells were transformed accordingly. The vectors are ready to be used for the heterologous expression of *Mf-folD* and *Mf-ftfL* in *E. coli* strains BL21 or Rosetta2 and subsequent His-tag purification for biochemical studies.

## METHODS

### DNA isolation, primer design and gene amplification

*M. fumariolicum* SolV was grown as methane-limited continuous culture as described before (Schmitz et al., 2020), except that a small 400 mL chemostat was used. A 5 mL sample was taken from the chemostat and centrifuged at 5000 × *g* at 4 °C for 5 min. The supernatant was removed and the pelleted cells were used for DNA isolation using the DNeasy PowerSoil Kit (Qiagen, Hilden, Germany) according to the manufacturer’s instructions. Forward and reverse primers were designed (**Table 1**) and subsequently produced by Biolegio (Nijmegen, The Netherlands) in order to amplify Mf-*folD* and Mf-*ftfL* from the template DNA. Mf-*folD* and *Mf-ftfL* were amplified using the following PCR mixture: 15 μL 2x Phusion High-Fidelity PCR Master Mix (Thermo Fisher Scientific, Waltham, MA, USA) was mixed with 0.5 μL template DNA and 13.3 μL Milli-Q water (MQ). To amplify *Mf-folD*, 0.6 μL 4 μM FolD_FW and 0.6 μL 4 μM FolD_REV were added to the PCR mixture. To amplify *Mf-ftfL*, 0.6 μL 4 μM FtfL_FW and 0.6 μL 4 μM FtfL_REV were added. Amplification was initiated by heating in a thermocycler at 98 °C for 30 sec to separate the template DNA into two single strands. Subsequently, 30 cycles were performed at 98 °C for 10 sec, 60 °C for 20 sec and 72 °C for 30 sec, for denaturation of double-stranded DNA, annealing of the primers and subsequent elongation, respectively. Hereafter, the samples were heated at 72 °C for 1 min as final elongation step. The PCR products were purified using the GeneJET PCR Purification Kit (Thermo Fisher Scientific), using 50 μL MQ instead of elution solution. The quality and concentrations of *Mf-folD* (24.5 ng *·*μL^-1^) and *Mf-ftfL* (31.3 ng *·*μL^-1^) were determined using a NanoDrop 1000 spectrophotometer (Thermo Fisher Scientific).

**Table 1:**
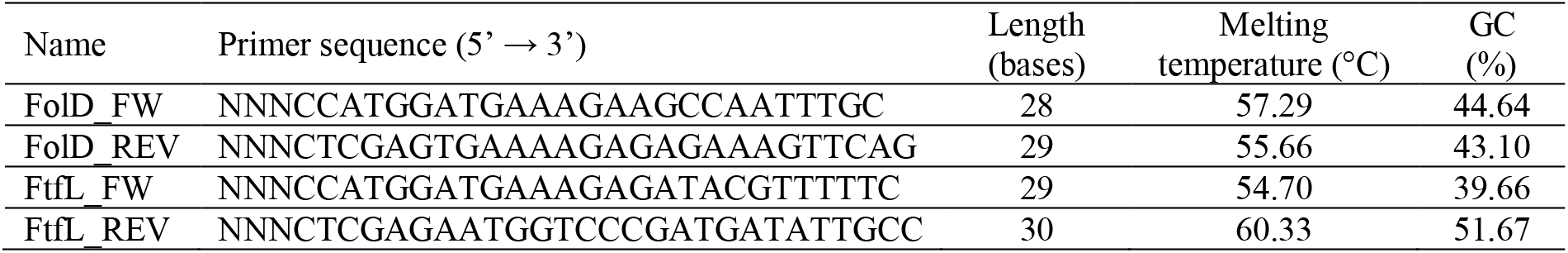
Forward and reverse primers for the amplification of *Mf-folD* and *Mf-ftfL*.

### Vector construction, restriction and ligation

To introduce *Mf-folD* and *Mf-ftfL* into a host cell, a pET30a(+) vector was constructed. This vector contains a kanamycin resistance gene and allows the fusion of a hexahistidine-tag at the C-terminus or N-terminus for His-tag purification after expression. An *E. coli* strain containing a pET30a(+) vector was taken from a glycerol stock stored at −80 °C. The cells were incubated in sterile liquid LB medium containing 50 μg · mL^-1^ kanamycin at 37 °C overnight at 250 rpm. The pET30a(+) vectors were harvested using the GeneJET Plasmid Miniprep Kit (Thermo Fisher Scientific) at a concentration of 96.7 ng *·*μL^-1^. Restriction of the PCR products *Mf-folD* and *Mf-ftfL* was performed by mixing 1 μL 10x FastDigest Buffer (Thermo Fisher Scientific) with 3 μL MQ, 0.5 μL 10 U *·*μL^-1^ NcoI (Thermo Fisher Scientific), 0.5 μL 10 U *·*μL^-1^ XhoI (Thermo Fisher Scientific) and 5 μL *Mf-folD* (24.5 ng *·*μL^-1^) or 5 μL *Mf-ftfL* (31.3 ng *·*μL^-1^). Restriction of the pET30a(+) vector was conducted similarly but with 3 μL vector (96.7 ng *·*μL^-1^) and 5 μL MQ instead. Samples were incubated at 37 °C for 30 min and subsequently the restriction enzymes were denatured by incubation at 80 °C for 2 min. To ligate Mf-*folD* and *Mf-ftfL* into the vector, 1 μL 10x T4 DNA Ligase Buffer (Thermo Fisher Scientific), 2.5 μL MQ, 1 μL T4 DNA ligase (Thermo Fisher Scientific), 0.5 μL restricted pET30a(+) vector (14.5 ng) and either 5 μL restricted *Mf-folD* (61.3 ng) or 5 μL restricted *Mf-ftfL* (78.3 ng) were mixed. The T4 DNA ligase was added last. As negative control, a mixture of the restricted vector without *Mf-folD* or *Mf-ftfL* was used. Samples were incubated at RT for 3 hours.

### Transformation

To transform the vectors into host cells, 2 μL of the ligation mixtures (either containing empty vectors, vectors with Mf-*folD* inserted or vectors with Mf-*ftfL* inserted) were added to Eppendorf tubes containing 50 μL chemically competent *E. coli* XL1-Blue cells. An additional Eppendorf tube containing 50 μL *E. coli* XL1-Blue cells was used and 2 μL MQ was added as control. The mixtures were put on ice for 5 min. Subsequently, the Eppendorf tubes were placed in a water bath at 42 °C for 90 sec. Hereafter, the samples were again incubated on ice for 5 min after which the cells were centrifuged for 1 min to pellet the cells. The supernatant was removed and cells were resuspended in 200 μL MQ and plated immediately. To observe whether transformation of the pET30a(+) vector was successful, the *E. coli* XL1-Blue cells were subsequently spread out with a Drigalski spatula on an LB plate containing 50 μg · mL^-1^ kanamycin. The plates were incubated overnight at 37 °C.

To observe whether the *E. coli* XL1-Blue cells were successfully transformed with vectors containing Mf-*folD*, vectors containing *Mf-ftfL* and the empty vectors, colony PCR was performed. A PCR mixture was prepared by mixing 75 μL 2x PerfeCTa SYBR Green FastMix (Quanta Bio, Beverly, MA, USA), 6 μL forward pET30a(+) primer, 6 μL reverse pET30a(+) primer and 63 μL MQ. Colonies of the four plates were labelled, picked with a sterile pipet tip and mixed with 10 μL of the mixture. Amplification of the vectors was initiated by heating in a thermocycler at 95 °C for 3 min to separate the DNA into two single strands. Subsequently, 30 cycles were performed at 95 °C for 30 sec, 58 °C for 30 sec and 72 °C for 2 min, for denaturation, annealing and elongation, respectively. Hereafter, the samples were heated at 72 °C for 10 min as a final elongation step. To sequence inserts of the pET30a(+) vectors to confirm successful insertion of *Mf-folD* and *Mf-ftfL*, the colonies obtained from the plates were placed in liquid LB medium containing 50 μg · mL^-1^ kanamycin and incubated overnight at 37 °C. The vectors were subsequently harvested with the GeneJET Plasmid Miniprep Kit (Thermo Fisher Scientific). Finally, two colonies of the *Mf-folD* plate (24.3 and 27.1 ng *·*μL^-1^) and two colonies of the *Mf-ftfL* plate (18.2 and 28.5 ng *·*μL^-1^) were picked and sequenced by BaseClear (Leiden, The Netherlands) using T7-forward and T7-reverse primers.

### Gel electrophoresis

To visualize PCR products, gels were prepared by mixing 0.29 g agarose with 30 mL 1% sodium borate buffer. The mixture was dissolved by intermittent heating and stirring. Hereafter, the mixture was cooled for 3 min and 3 μL EtBr was added and polymerized for 20 min. 3.5 μL GeneRuler 1kb Plus DNA Ladder (Thermo Fisher Scientific) was used as molecular marker to assess the size of the PCR products. 5 μL PCR sample with 1 μL loading dye were used and run for 30 min at 70 V.

### Amino acid sequence comparisons

The amino acid sequences of FolD of *M. extorquens* strain CM4 (WP_012606308.1), FolD of *E. coli* (WP_187226985.1) and FtfL of *M. extorquens* AM1 (WP_003606333.1) were retrieved from GenBank. FolD and FtfL of *M. extorquens* were used for comparison since methylotrophy has been well studied in this model organism (Studer et al., 2002; Marx et al., 2003; Kim et al., 2020). FolD of *E. coli* was used for comparison since this enzyme was purified and biochemically characterized, whereas FolD of *M. extorquens* has not been isolated (D’ Ari and Rabinowitz, 1991). InterPro was used to predict protein families (Blum et al., 2021).

## RESULTS AND DISCUSSION

*Mf-folD* (Mfumv2_1033) encodes a cytoplasmic protein of 297 amino acids that was shown to be expressed when cells were grown on methane (Mohammadi et al., 2017). The gene is annotated as “methylenetetrahydrofolate dehydrogenase (NADP^+^)/Methenyltetrahydrofolate cyclohydrolase (folD)” and InterPro categorizes it as a member of the protein family “tetrahydrofolate dehydrogenase/cyclohydrolase (IPR000672)”. The amino acid sequence of Mf-FolD is 42% identical (63% positives; E = 7^-73^) compared to FolD of *M. extorquens* strain CM4 and 43% identical (63% positives; E = 7^-76^) compared to FolD of *E. coli*. The purified enzyme from *E. coli* was shown to convert methylene-tetrahydrofolate (CH_2_-H_4_F) to formyl-tetrahydrofolate (CHO-H_4_F) (D’Ari and Rabinowitz, 1991). *Mf-ftfL* (Mfumv2_2082) encodes a cytoplasmic protein of 563 amino acids that is expressed at a similar quantity as *Mf-folD*.The gene is annotated as “formate—tetrahydrofolate ligase” and InterPro categorizes it as a member of the protein family “Formate-tetrahydrofolate ligase, FTHFS (IPR000559)”. The amino acid sequence of Mf-FtfL is 50% identical (67% positives; E = 0.0) compared to FtfL of *M. extorquens* AM1 of which the purified enzyme was shown to convert CHO-H4F to formate and vice versa (Marx et al., 2003; Kim et al., 2020). Amino acid sequences most closely related to FolD and FtfL of verrucomicrobial methanotrophs are found in the phyla Verrucomicrobia and Proteobacteria, respectively.

Whereas many methylotrophs possess multiple pathways to modulate the dissimilation and assimilation of one-carbon compounds, genomic analyses of verrucomicrobial methanotrophs suggest a relatively simplistic mode of methylotrophy (Chistoserdova, 2011; Schmitz et al., 2021). The formaldehyde oxidation pathway that utilizes the coenzyme tetrahydromethanopterin (H_4_MPT) as C1-carrier has been well studied in *M. extorquens* and is found in a range of methylotrophs (Chistoserdova et al., 1998; Vorholt et al., 1999; Vorholt 2002; Chistoserdova et al., 2009; Chistoserdova, 2011). In this pathway, the formaldehyde-activating enzyme (Fae) catalyses the condensation of H_4_MPT with formaldehyde to eventually synthesize formate via several intermediates (Vorholt et al., 2000; Crowther et al., 2008). However, genes involved in the H_4_MPT-dependent pathway are not encoded by verrucomicrobial methanotrophs and also Fae is absent. In addition, several organisms oxidize formaldehyde through a glutathione-dependent pathway (Vorholt, 2002), which is also not found in verrucomicrobial methanotrophs. Finally, many methylotrophs convert one-carbon compounds by using tetrahydrofolate (H_4_F) as C1-carrier (Chistoserdova, 2011). A pathway to synthesize H_4_F seems to be present in verrucomicrobial methanotrophs (Hou et al., 2008; Khadem et al., 2012; Schmitz et al., 2021). In *M. extorquens* AM1, this pathway is thought to be primarily used to generate CH_2_-H_4_F to feed the serine cycle (Crowther et al., 2008). As such, formate generated through the H_4_MPT-dependent pathway is converted by FtfL to CHO-H_4_F, which is converted to CH-H_4_F by the methenyl-H_4_F cyclohydrolase (Fch) and subsequently to CH_2_-H_4_F by the methylene-H_4_F dehydrogenase (MtdA) (Pomper et al., 1999; Marx and Lidstrom, 2004; Crowther et al., 2008).

Enzymes involved in the H_4_F-dependent pathway are reversible and the pathway can therefore work in an oxidative and reductive direction (Studer et al., 2002). Whereas FtfL is present in all verrucomicrobial methanotrophs, the genes encoding Fch and MtdA are absent (Chistoserdova 2011; Schmitz et al., 2021). The enzyme FolD, however, was shown to be a bifunctional enzyme that can catalyse the conversion of CH_2_-H_4_F to CHO-H_4_F or vice versa (D’Ari and Rabinowitz, 1991; Vannelli et al., 1999; Studer et al., 2002). Consequently, verrucomicrobial methanotrophs might use FolD and FtfL to catalyse the oxidative conversion of formaldehyde to formate, rendering ATP and NAD(P)H (**Figure 1**). A complicating factor in employing the H4F-dependent pathway in an oxidative direction could be the formation of CH_2_-H_4_F (Vorholt et al., 1999; Marx et al., 2005; Crowther et al., 2008). Although formaldehyde binds spontaneously to H_4_F, this condensation was shown to be too slow for *E. coli* to grow efficiently on methanol (Kallen and Jencks, 1966; He et al., 2020). On the other hand, no enzyme is found in the model methylotrophic microorganism *M. extorquens* that could enhance this reaction (Vorholt et al., 1999). Hence, an enzyme might not be necessary for the formation of CH_2_-H_4_F, or a yet to be discovered enzyme is involved.

**Figure 1:**
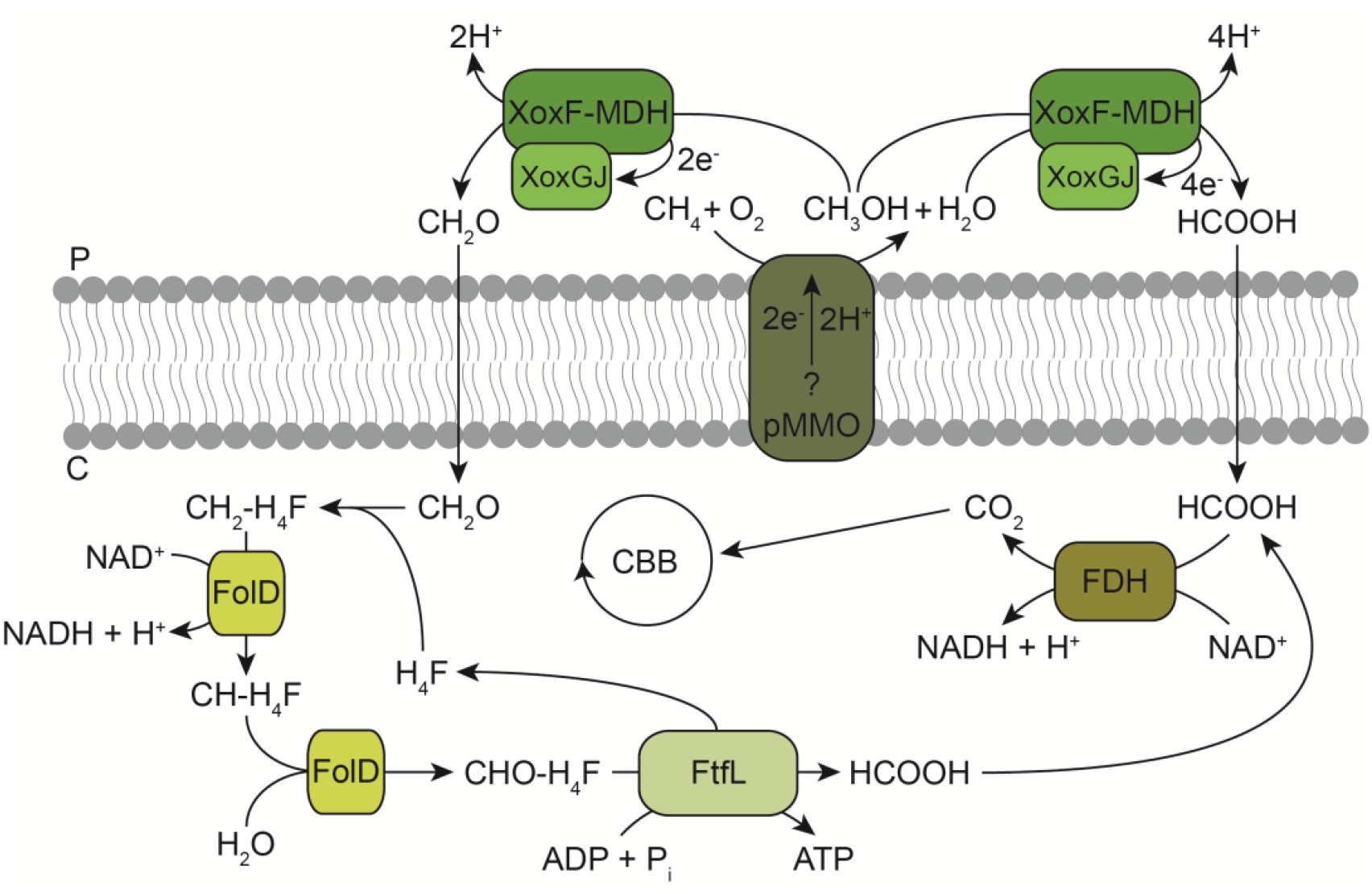
An alternative pathway for formaldehyde oxidation in verrucomicrobial methanotrophs. pMMO oxidizes methane to methanol (CH_3_OH), while an unknown electron donor is oxidized. The lanthanide-dependent XoxF methanol dehydrogenase (MDH) could subsequently oxidize methanol to either formate (HCOOH) or formaldehyde (CH_2_O), while donating electrons to its redox partner XoxGJ. Formate diffuses into the cytoplasm and is converted to CO_2_ by an NAD^+^-dependent formate dehydrogenase (FDH). CO_2_ is fixed into biomass via the Calvin-Benson-Bassham (CBB) cycle. Alternatively, formaldehyde could bind to tetrahydrofolate (H_4_F) spontaneously to form methylene-tetrahydrofolate (CH_2_-H_4_F). The enzyme FolD converts CH_2_-H_4_F to methenyl-tetrahydrofolate (CH-H_4_F), which is subsequently converted to formyl-tetrahydrofolate (CHO-H_4_F). This product is then converted to H_4_F and formate, while producing ATP. Adapted from Schmitz et al. (2021).

The reactions catalysed by FolD and FtfL have been shown before in various microorganisms, but not in verrucomicrobial methanotrophs (Goenrich et al., 2002; Marx et al., 2003). The utilization of FtfL and FolD in a reductive direction would be illogical in verrucomicrobial methanotrophs, since these microbes do not possess a complete serine cycle and because they were shown to use the CBB cycle for CO_2_ fixation (Khadem et al., 2011; Van Teeseling al., 2014). Alternatively, FolD and FtfL are dedicated to other or multiple functions in the cell. Indeed, CHO-H_4_F produced by FolD is a precursor for purine biosynthesis (He et al., 2020). The necessity of the H_4_F-dependent pathway for formaldehyde oxidation to formate in verrucomicrobial methanotrophs is determined by the product formation of XoxF *in vivo* (**Figure 1**). If XoxF produces formate in the periplasm, the H_4_F-dependent pathway would be largely redundant and formate could subsequently be converted to CO_2_ in the cytoplasm by the formate dehydrogenase (Keltjens et al., 2014). Nevertheless, in this scenario the H_4_F-dependent pathway might still be useful in case of leakage of toxic formaldehyde into the cytoplasm (Vorholt et al., 2000).

To enable heterologous expression of Mf-FolD and Mf-FtfL in *E. coli*, the genes encoding these enzymes were amplified using the designed primers. Subsequently, the PCR products were restricted and ligated into pET30a(+) vectors. Gel electrophoresis of the products obtained through colony PCR revealed successful transformation of the *E. coli* XL1-Blue cells with an empty vector or a vector in which Mf-*folD* or Mf-*ftfL* was ligated (data not shown). Indeed, a colony putatively containing a pET30a(+) vector with the reverse complement DNA sequence of Mf-*folD* inserted possesses an insert that maps 100% to Mf-*folD* (**Supplementary Table S1**). In addition, a colony putatively containing a pET30a(+) vector with Mf-*ftfL* inserted possesses an insert that maps almost 100% to Mf-*ftfL* (**Supplementary Table S1**). The DNA sequence ligated into the pET30a(+) vector possesses one base difference compared to the sequence of Mf-*ftfL* in the genome. The gene Mf-*ftfL* in the genome starts with the codon GTG. This codon encodes the amino acid valine when present inside a sequence, but when present as the first codon it functions as alternative start codon and is translated to a methionine residue (Lobanov et al., 2010). The Mf-*ftfL* forward primer FtfL_FW was designed in such a way that the gene would start with ATG, which is also translated into methionine. The start codon ATG was used because it is more common in *E. coli* and explains the single base difference between Mf-*ftfL* inserted in the vector and Mf-*ftfL* in the genome of *M. fumariolicum* SolV (**Supplementary Table S1**). Both the Mf-*folD* and the Mf-*ftfL* DNA sequences in the vector have additional nucleotides attached that are translated into the hexahistidine-tag LEHHHHHH for His-tag purification (**Table 2**).

**Table 2:**
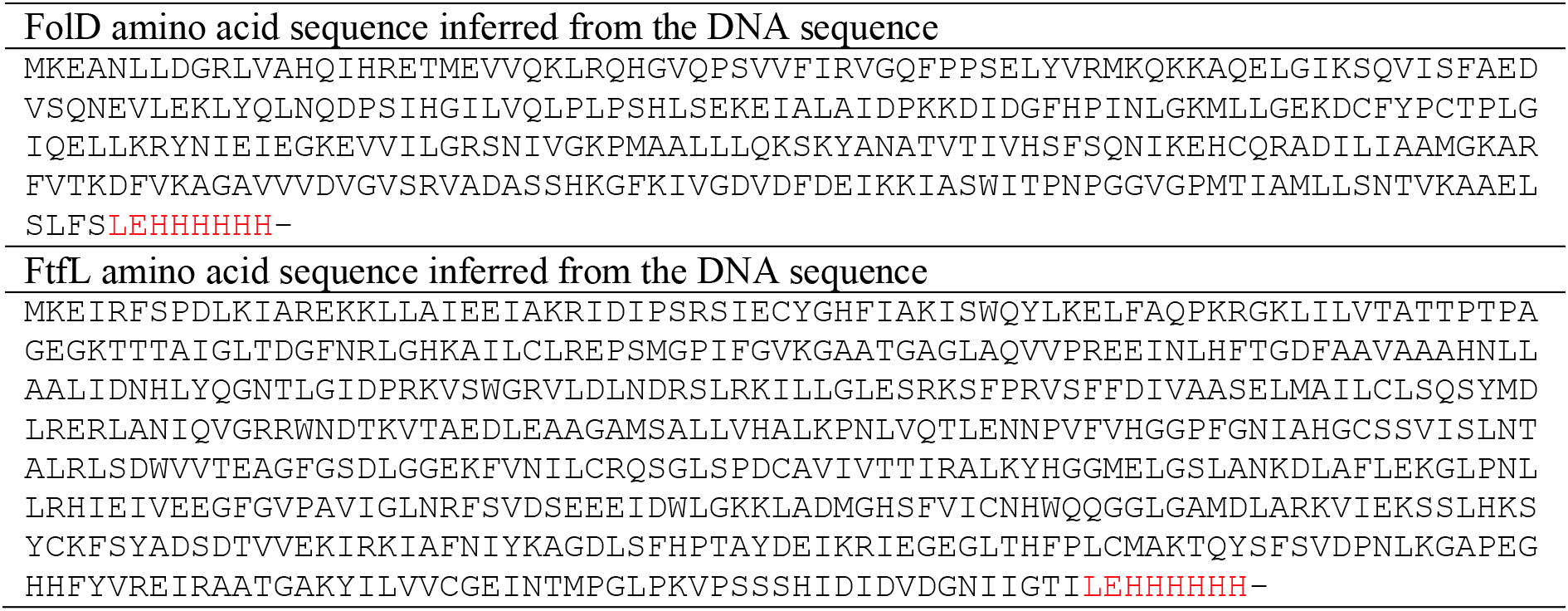
Amino acid sequences that can be produced from Mf-*folD* and Mf-*ftfL* present in the pET30a(+) vectors.

The production of formate by purified XoxF (type XoxF2) of *M. fumariolicum* SolV was questioned by the observation that a related XoxF-type MDH (type XoxF5) of *M. extorquens* AM1 was shown to produce formaldehyde *in vivo* (Good et al., 2019). XoxF2-type MDHs are only found in verrucomicrobial methanotrophs and *‘Candidatus* Methylomirabilis oxyfera’, whereas XoxF5-type MDHs are only found in Proteobacteria (Chistoserdova 2011; Keltjens et al., 2014). Interestingly, XoxF2 of *M. fumariolicum* SolV has an almost ten times higher affinity for formaldehyde than XoxF5 of *M. extorquens* AM1 (Keltjens et al., 2014). In addition, in *M. extorquens* AM1 the H4MPT-dependent pathway is dedicated to the oxidation of formaldehyde to formate, whereas in verrucomicrobial methanotrophs it is not (Good et al., 2019). Altogether, the product formed by XoxF might depend on the specific type of XoxF, on the presence of a pathway dedicated to the oxidation of formaldehyde to formate and on the intermediate used in carbon assimilation.

In conclusion, the primers designed and transformation performed in this study can be used to purify Mf-FolD and Mf-FtfL to validate the hypothesized function. In addition, metabolic flux analyses could be used to decipher how the alternative tetrahydrofolate pathway for formaldehyde oxidation could be employed by verrucomicrobial methanotrophs (Marx et al., 2005). Lastly, ultimate proof would be to knockout Mf-*folD* and Mf-*ftfL* when a genetic system becomes available to observe whether growth on methane and methanol is still possible.

## AUTHOR CONTRIBUTIONS

RAS, KAJP and HJMOdC designed the project and experiments. RAS and KAJP conducted the experiments and data analyses. RAS and HJMOdC wrote the manuscript. HJMOdC supervised the research.

## ACKNOWLEDGEMENTS

RAS and HJMOdC were supported by the European Research Council (ERC Advanced Grant project VOLCANO 669371) and KAJP was supported through the Gravitation Grant Netherlands Earth System Science Centre (grant number 024.002.001).

## REFERENCES

Blum M, Chang HY, Chuguransky S et al. The InterPro protein families and domains database: 20 years on. Nucleic Acids Res 2021;49:D344–D354.

Chistoserdova L, Vorholt JA, Thauer RK, Lidstrom ME. C1 transfer enzymes and coenzymes linking methylotrophic bacteria and methanogenic Archaea. Science 1998;281:99–102.

Chistoserdova L, Kalyuzhnaya MG, Lidstrom ME. The expanding world of methylotrophic metabolism. Annu Rev Microbiol 2009;63:477–499.

Chistoserdova L. Modularity of methylotrophy, revisited. Environ Microbiol 2011;13:2603–2622.

Chistoserdova L, Kalyuzhnaya MG. Current trends in methylotrophy. Trends Microbiol 2018;26:703–714.

Crowther GJ, Kosály G, Lidstrom ME. Formate as the main branch point for methylotrophic metabolism in *Methylobacterium extorquens* AM1. J Bacteriol 2008;190:5057–5062.

D’Ari L, Rabinowitz JC. Purification, characterization, cloning, and amino acid sequence of the bifunctional enzyme 5,10-methylenetetrahydrofolate dehydrogenase/5,10-methenyltetrahydrofolate cyclohydrolase from *Escherichia coli*. J Biol Chem 1991;266:23953–23958.

Goenrich M, Bursy J, Hübner E, Linder D, Schwartz AC, Vorholt JA. Purification and characterization of the methylene tetrahydromethanopterin dehydrogenase MtdB and the methylene tetrahydrofolate dehydrogenase FolD from *Hyphomicrobium zavarzinii* ZV580. Arch Microbiol 2002;177:299–303.

Good NM, Moore RS, Suriano CJ, Martinez-Gomez NC. Contrasting *in vitro* and *in vivo* methanol oxidation activities of lanthanide-dependent alcohol dehydrogenases XoxF1 and ExaF from *Methylobacterium extorquens* AM1. Sci Rep 2019;9:4248.

Hanson RS, Hanson TE. Methanotrophic bacteria. Microbiol Rev 1996;60:439–471.

He H, Noor E, Ramos-Parra PA, García-Valencia LE, Patterson JA, Díaz de la Garza RI et al. *In vivo* rate of formaldehyde condensation with tetrahydrofolate. Metabolites 2020;10:65.

Hou S, Makarova KS, Saw JHW, Senin P, Ly BV, Zhou Z et al. Complete genome sequence of the extremely acidophilic methanotroph isolate V4, *Methylacidiphilum infernorum*, a representative of the bacterial phylum Verrucomicrobia. Biol Direct 2008;3:26.

Kallen RG, Jencks WP. The mechanism of the condensation of formaldehyde with tetrahydrofolic acid. J Biol Chem 1966;241:5851–5863.

Keltjens JT, Pol A, Reimann J, Op den Camp HJM. PQQ-dependent methanol dehydrogenases: rare-earth elements make a difference. Appl Microbiol Biotechnol 2014;98:6163–6183.

Khadem AF, Pol A, Wieczorek AS, Mohammadi SS, Francoijs K-J, Stunnenberg HG et al. Autotrophic methanotrophy in Verrucomicrobia: *Methylacidiphilum fumariolicum* SolV uses the Calvin-Benson-Bassham cycle for carbon dioxide fixation. J Bacteriol 2011;193:4438–4446.

Khadem AF, Pol A, Wieczorek AS, Jetten MSM, Op den Camp HJM. Metabolic regulation of “*Ca. Methylacidiphilum fumariolicum”* SolV cells grown under different nitrogen and oxygen limitations. Front Microbiol 2012;3:266.

Kim S, Lee SH, Seo H, Kim KJ. Biochemical properties and crystal structure of formate-tetrahydrofolate ligase from *Methylobacterium extorquens* CM4. Biochem Biophys Res Commun 2020;528:426–431.

Lobanov AV, Turanov AA, Hatfield DL, Gladyshev VN. Dual functions of codons in the genetic code. Crit Rev Biochem Mol Biol 2010;45:257–265.

Marx CJ, Laukel M, Vorholt JA. Purification of the formatetetrahydrofolate ligase from *Methylobacterium extorquens* AM1 and demonstration of its requirement for methylotrophic growth. J Bacteriol 2003;185:7169–7175.

Marx CJ, Lidstrom ME. Development of an insertional expression vector system for *Methylobacterium extorquens* AM1 and generation of null mutants lacking mtdA and/or fch. Microbiology 2004;150:9–19.

Marx CJ, van Dien SJ, Lidstrom ME. Flux analysis uncovers key role of functional redundancy in formaldehyde metabolism. PLoS Biol 2005;3:e16.

Mohammadi SS, Pol A, van Alen TA, Jetten MSM, Op den Camp HJM. *Methylacidiphilum fumariolicum* SolV, a thermoacidophilic ‘Knallgas’ methanotroph with both an oxygen-sensitive and -insensitive hydrogenase. ISME J 2017;11:945–958.

Op den Camp HJM, Islam T, Stott MB, Harhangi HR, Hynes A, Schouten S et al. Environmental, genomic and taxonomic perspectives on methanotrophic Verrucomicrobia. Environ Microbiol Rep 2009;1:293–306.

Picone N, Op den Camp HJM. Role of rare earth elements in methanol oxidation. Curr Opin Chem Biol 2019;49:39–44.

Picone N, Blom P, Hogendoorn C, Frank J, van Alen T, Pol A et al. Metagenome assembled genome of a novel verrucomicrobial methanotroph from Pantelleria Island. Front Microbiol 2021;12:666929.

Pol A, Barends TRM, Dietl A, Khadem AF, Eygensteyn J, Jetten MSM et al. Rare earth metals are essential for methanotrophic life in volcanic mudpots. Environ Microbiol 2014;16:255–264.

Pomper BK, Vorholt JA, Chistoserdova L, Lidstrom ME, Thauer RK. A methenyl tetrahydromethanopterin cyclohydrolase and a methenyl tetrahydrofolate cyclohydrolase in *Methylobacterium extorquens* AM1. Eur J Biochem 1999;261:475–480.

Ross MO, Rosenzweig AC. A tale of two methane monooxygenases. J Biol Inorg Chem 2017;22:307–319.

Schmitz RA, Pol A, Mohammadi SS, Hogendoorn C, van Gelder AH, Jetten MSM et al. The thermoacidophilic methanotroph *Methylacidiphilum fumariolicum* SolV oxidizes subatmospheric H2 with a high-affinity, membrane-associated [NiFe] hydrogenase. ISME J 2020;14: 1223–1232.

Schmitz RA, Peeters SH, Versantvoort W, Picone N, Pol A, Jetten MSM et al. Verrucomicrobial methanotrophs: ecophysiology of metabolically versatile acidophiles. FEMS Microbiol Rev 2021;fuab007.

Studer A, McAnulla C, Buchele R, Leisinger T, Vuilleumier S. Chloromethane-induced genes define a third C_1_ utilization pathway in *Methylobacterium chloromethanicum* CM4. J Bacteriol 2002;184:3476–3484.

van Teeseling MC, Pol A, Harhangi HR, van der Zwart S, Jetten MSM, Op den Camp HJM et al. Expanding the verrucomicrobial methanotrophic world: description of three novel species of *Methylacidimicrobium* gen. nov. Appl Environ Microbiol 2014;80:6782–6791.

Vannelli T, Messmer M, Studer A, Vuilleumier S, Leisinger T. A corrinoid-dependent catabolic pathway for growth of a *Methylobacterium* strain with chloromethane. Proc Natl Acad Sci U S A 1999;96:4615–4620.

Vorholt JA, Chistoserdova L, Stolyar SM, Thauer RK, Lidstrom ME. Distribution of tetrahydromethanopterin-dependent enzymes in methylotrophic bacteria and phylogeny of methenyl tetrahydromethanopterin cyclohydrolases. J Bacteriol 1999;181:5750–5757.

Vorholt JA, Marx CJ, Lidstrom ME, Thauer RK. Novel formaldehyde-activating enzyme in *Methylobacterium extorquens* AM1 required for growth on methanol. J Bacteriol 2000;182: 6645–6650.

Vorholt JA. Cofactor-dependent pathways of formaldehyde oxidation in methylotrophic bacteria. Arch Microbiol 2002;178:239–249.

