## Supplementary Table 1 for "An alternative tetrahydrofolate pathway for formaldehyde oxidation in verrucomicrobial methanotrophs: Primer design for *folD* and *ftfL* and transformation of *E. coli*" (from another version of this preprint: version 1, 2022-08-24)

### SUPPLEMENTARY INFORMATION

**Supplementary Table S1**: DNA sequences Mf-*folD* or Mf-*ftfL* present in the pET30a(+) vectors. Red bases indicate C-terminal sequences encoding His-tags. Bases with yellow background indicate stop codons. The base with a green background indicates a base difference compared to the genome if *Methylacidiphilum fumariolicum* SolV.

| Reverse complement *folD* DNA sequence in the pET30(+) vector |
|---|
| tcagtggtggtggtggtggtgctcgagTGAAAAGAGAGAAAGTTCAGCGGCTTTAACGGTATTGGATAACAGCATTGCTAT |
| GGTCATGGGACCAACTCCTCCTGGATTTGGAGTAATCCAGGAGGCTATTTTTTTTATTTCATCAAAATCAACATCTCCAAC |
| AATCTTAAATCCCTTATGTGAGGAAGCATCAGCAACCCTGCTAACTCCAACATCAACAACCACTGCGCCAGCTTTGACAAA |
| ATCTTTTGTTACAAACCGAGCTTTACCCATTGCGGCTATCAGGATATCGGCTCTCTGACAATGTTCTTTGATGTTTTGACT |
| GAAGGAATGGACAATAGTCACCGTTGCATTCGCATACTTCGATTTTTGGAGTAGTAGAGCTGCCATTGGTTTCCCAACAAT |
| ATTGCTTCTTCCAAGGATAACCACTTCTTTTCCTTCGATCTCGATGTTGTAGCGCTTCAACAGTTCTTGAATACCCAATGG |
| AGTGCACGGATAAAAGCAGTCTTTTTCTCCAAGGAGCATTTTCCCAAGGTTTATAGGATGAAATCCATCAATATCCTTTTT |
| GGGATCAATAGCAAGAGCTATTTCTTTCTCAGATAGATGGCTGGGAAGTGGAAGTTGAACCAGTATACCGTGGATGGAAGG |
| ATCTTGATTTAATTGATAGAGTTTCTCTAAGACTTCGTTTTGTGATACATCCTCAGCAAAAGAAATGACTTGGGATTTGAT |
| CCCAAGTTCTTGAGCTTTCTTCTGTTTCATCCTAACATAAAGTTCAGATGGTGGGAACTGTCCAACTCTTATAAAAACGAC |
| AGAAGGTTGAACTCCATGTTGTCGCAACTTCTGAACAACTTCCATTGTTTCTCTATGGATCTGATGTGCCACAAGCCTACC |
| GTCCAGCAAATTGGCTTCTTTCAT |
| *ftfL* DNA sequence in the pET30(+) vector |
| ATGAAAGAGATACGTTTTTCTCCTGATTTAAAGATAGCACGCGAAAAAAAACTTTTAGCCATAGAGGAAATAGCAAAAAGA |
| ATTGATATTCCTTCAAGATCCATTGAATGTTACGGACATTTTATTGCCAAAATATCATGGCAATATTTAAAAGAGCTTTTT |
| GCTCAGCCTAAGCGAGGTAAGCTCATTTTGGTGACAGCTACGACACCTACTCCAGCTGGGGAAGGAAAAACGACGACGGCG |
| ATTGGCCTCACTGATGGATTCAATCGATTAGGTCATAAAGCCATTCTTTGTTTGAGAGAGCCATCGATGGGCCCCATTTTT |
| GGAGTGAAAGGTGCGGCTACCGGGGCGGGTCTGGCACAAGTTGTTCCAAGAGAAGAGATAAATCTTCATTTTACTGGTGAT |
| TTTGCCGCAGTGGCGGCCGCTCATAATCTTCTGGCTGCCTTGATAGATAACCATCTTTATCAGGGTAATACCTTAGGTATT |
| GATCCAAGAAAGGTAAGCTGGGGAAGGGTATTGGATTTAAATGACAGGTCTTTGCGAAAAATCCTTCTGGGGTTAGAAAGT |
| AGAAAATCTTTTCCCAGAGTCTCTTTTTTTGACATCGTAGCAGCATCTGAATTGATGGCCATATTGTGCCTTTCGCAATCC |
| TACATGGATTTGAGGGAAAGGTTAGCCAACATCCAGGTTGGCCGCAGATGGAACGATACAAAGGTTACAGCCGAAGATCTT |
| GAGGCTGCTGGGGCTATGTCGGCCCTTTTAGTGCATGCTCTCAAACCCAATCTTGTCCAAACTTTGGAAAATAATCCAGTT |
| TTTGTGCATGGGGGACCTTTTGGCAATATTGCTCATGGATGCAGTTCGGTGATATCATTGAATACGGCCTTGAGGCTTTCT |
| GATTGGGTGGTTACTGAAGCAGGATTTGGCAGTGATCTAGGAGGAGAAAAATTCGTTAACATCCTGTGCCGGCAGAGTGGT |
| TTAAGCCCAGACTGTGCGGTGATAGTCACGACTATTCGTGCTTTAAAGTATCATGGAGGGATGGAGTTGGGATCCCTGGCA |
| AATAAAGACTTAGCCTTTTTAGAAAAGGGACTGCCTAATCTTTTAAGACATATTGAGATTGTCGAAGAAGGATTTGGAGTA |
| CCTGCAGTCATTGGACTGAACCGTTTTAGCGTTGATAGTGAAGAGGAAATCGATTGGCTTGGCAAAAAACTTGCCGACATG |
| GGACATTCCTTTGTCATATGCAATCATTGGCAGCAAGGAGGGCTAGGGGCTATGGATCTTGCAAGGAAAGTGATCGAAAAA |
| AGTAGCCTCCATAAAAGTTATTGTAAGTTTAGCTATGCAGACTCGGACACTGTTGTCGAAAAGATTCGAAAGATCGCATTT |
| AATATTTATAAAGCTGGAGATCTTTCCTTTCATCCTACTGCTTATGATGAGATAAAAAGGATTGAAGGAGAGGGACTAACC |
| CATTTTCCCCTATGTATGGCTAAAACGCAATATTCATTTTCTGTTGATCCCAATCTAAAAGGGGCGCCTGAAGGACACCAT |
| TTTTATGTTAGAGAAATCAGAGCGGCTACTGGGGCTAAATACATCCTTGTTGTGTGTGGGGAGATAAATACGATGCCAGGC |
| TTGCCCAAAGTCCCTTCTTCCAGTCATATCGACATCGATGTGGATGGCAATATCATCGGGACCATTctcgagcaccaccac |
| caccaccactga |
